# Letters: Complex response of beta diversity to dispersal in meta-community models

**DOI:** 10.1101/2021.02.19.432043

**Authors:** Muyang Lu

## Abstract

Dispersal is one of the most important drivers of community assembly. The conventional belief that dispersal leads to biotic homogenization (lower beta diversity) has been recently challenged by an experiment conducted in nectar microbes (Vannette & Fukami, 2017), showing that dispersal could lead to community divergence. In this paper, I re-examined the relationship between beta diversity and local dispersal in a range of theoretical models: from the classic island biogeography model and meta-population model to a meta-community model that incorporates biotic interactions. I find that the emergence of hump-shaped beta diversity-dispersal relationship is closely related to local dispersal (rather than global dispersal), non-neutrality and biotic interactions. The results reveal rich metacommunity dynamics in relation to dispersal types and biotic interactions which might be overlooked in previous theoretical and empirical studies. The findings call for more realistic experimental manipulations on dispersals in future community assembly studies.

## Introduction

Understanding the bewildering diversity and complexity of natural ecosystems is a fundamental goal of ecology. Species co-occurrence, or beta diversity (i.e. compositional dissimilarity) across communities is at the heart of this pursuit. Why do some species co-occur together, and why are some communities so distinct from each other? These questions have fueled longstanding debates in the field: from Clements and Gleason’s arguments over climax community (Clements 1916; Gleason 1926), to Diamond and Simberloff’s debates over checkerboard distributions (Connor & Simberloff 1978, 1979; Connor *et al.* 2015; Diamond *et al.* 2015), to the recent discussions over the drivers of latitudinal beta diversity gradients (Kraft *et al.* 2011; Qian *et al.* 2013; Xu *et al.* 2015; Sreekar *et al.* 2018; Xing & He 2019). The mechanisms of beta diversity formations revealed during these debates further stimulated the development of recent statistical tools to disentangle the effect of niche and neutral processes for community assembles (Chase & Myers 2011; Mori *et al.* 2015; Ning *et al.* 2019).

Dispersal, one of the most fundamental processes in community assembly, has received great attention from community ecologists (Grainger & Gilbert 2016). The conclusions about the effect of dispersal on beta diversity are remarkably consistent throughout the literature: various theoretical work (Hubbell 2001; Mouquet & Loreau 2003; Lu *et al.* 2019) and empirical studies (Gianuca *et al.* 2017; Ron *et al.* 2018) have shown that increasing dispersal leads to community homogenization. Partly due to the intuition of the arguments, i.e. more exchange of species among communities naturally leads to more similar community compositions, and partly because of the abundance of evidence in favor of the finding, the view of ‘dispersal leads to homogenization’ has remained unchallenged until very recently. In a nectar microbial experiment, Vannette and Fukami (2017) demonstrated contrary to the conventional belief, increasing dispersal could lead to community divergence because of priority effects. Their study casts doubt on whether a universal dispersal-beta diversity relationship truly exists, and points to a key factor in generating an increasing beta diversity-dispersal relationship – that dispersal being highly stochastic.

There are two ways of modeling dispersal stochasticity in established ecological theories: dispersal can either be an island-type dispersal which models global and external dispersal with immigrants originate from a mainland species pool and arrive at different patches (islands) independently (MacArthur & Wilson 1967), or a meta-population type dispersal (Hanski 1998) which models internal and local dispersal with immigrants coming from other patches. In light of this distinction, the neutral theory falls into the island-type as immigrants are drawn from a fixed meta-community (Hubbell 2001); most microcosm experiments and meta-community models, in which dispersal is implemented by exchanging migrants among patches, fall into meta-population dispersals (Mouquet & Loreau 2003; Ron *et al.* 2018). In most natural ecosystems, the dispersal is a mixture of the two: forest plots will receive seeds generated within the plot and migrated from outside; archipelagos will intercept immigrants both from the mainland and nearby islands. Whether the two types of dispersal create different beta diversity-dispersal relationship when examined in realistic biological contexts is unclear, as most experimental and theoretical work eliminate much of the potentially important heterogeneity in a natural system (Grainger & Gilbert 2016). I hypothesize that the island-type (global) dispersal is more likely to lead to community homogeneity than the meta-population (local) type dispersal as shared, constant species pool provides less room for local variation in community compositional changes.

Another key driver of the increasing beta diversity-dispersal relationship in Vannette and Fukami’s work is biotic interactions. Different forms of biotic interactions could have different impact on the beta diversity-dispersal relationship. For example, in the context of competition-colonization trade-off (Tilman 1994; Hanski 2008), equally increasing the dispersal rates of all species would ultimately lead to the elimination of the inferior competitor and community homogenization, while preferentially increasing the dispersal rate of the inferior competitor might facilitate regional coexistence and local community divergence; In apparent competitions (Holt 1977; Bonsall & Hassell 1997), increasing dispersal among habitats could lead to community homogenization if the abundance of the dominant prey species is suppressed by the introduction of the predator species, or lead to community divergence if different prey species are eliminated in different habitats by the predator species. In a similar vein, positive biotic interactions such as facilitation could lead to community homogenization with increasing dispersal if the facilitator is positively associated with all other species, or lead to community divergence if facilitation is highly specialized. In summary, the relationship between beta diversity and dispersal is far from straightforward when biotic interactions come into play. Other factors such as environmental heterogeneity (Gianuca *et al.* 2017) and habitat connectivity (Hewitt *et al.* 2005) further complicate the relationship between beta diversity and dispersal by modifying competition outcomes or creating more contingency in community assembly. A thorough understanding of the beta diversity-dispersal relationship hence necessitates a bottom-up theoretical investigation to disentangle the effects of different processes.

To progress toward this goal, I built from the classic island biogeography model (MacArthur & Wilson 1967) and meta-population model (Hanski 1998), and later combined them into a meta-community model. The purpose of this study is to illustrate the complexity of the beta diversity-dispersal relationship when different types of dispersal (local vs. global) and biotic interactions are simultaneously considered and provide guidance for further empirical work.

## Material and methods

I start with simple two-species occupancy models to demonstrate the difference between the island biogeography model and the meta-population model in beta diversity – dispersal relationship. I then extended the investigation to a spatially explicit meta-community model with more species and more islands using a discrete Markov model (see Cazelles *et al.* 2016). I used Jaccard dissimilarity as a metric for beta-diversity to better link the results to my previous work (Lu *et al.* 2019).

### The two-species, two-islands model

The formulation of my model is built on the island biogeography theory as in my previous paper (Lu *et al.* 2019). Consider two identical islands where the occurrence probability of species 1 is *p* and the occurrence probability of species 2 is *q*. By assuming that the species’ occurrences are independent between species and between islands, I can calculate the occurrence probability as well as its associated beta diversity (Supplement, Table S1). For example, the probability of having the two-islands meta-community in a state where species 1 is present on island 1 but absent on island 2, and species 2 is present on island 2 but absent on island 2, is *p*(1-*p*)*q*(1-*q*). The Jaccard dissimilarity between the two islands in this example is 1 since no species is shared between islands. The expected Jaccard dissimilarity for the two species - two islands model is calculated by summing the probability of each state of the two-island metacommunity and its corresponding Jaccard dissimilarity, which, after excluding double-absence state (Jaccard index is undefined at double-absence state; Anderson *et al.* 2011), is:

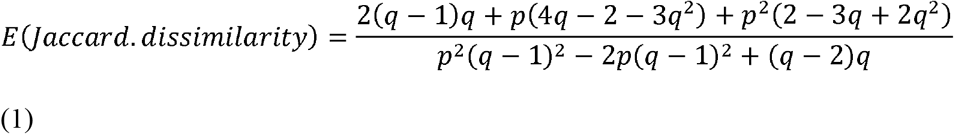

### The island biogeography model

In this model, immigrant species may only arrive from the mainland species pool (MacArthur & Wilson 1967). The occurrence probability of a species on an island is a function of extinction rate *e* and colonization rate *c*:

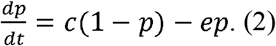

The equilibrium occurrence probability is:

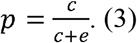

Assuming equivalence between two species, and substituting

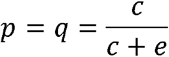

into equation (1), the expected Jaccard dissimilarity becomes:

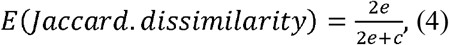

which is the same result from what I derived from the species-neutral island biogeography model in a previous paper (Lu et al. 2019).

For the non-neutral case, let

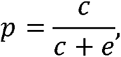

and

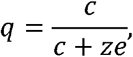

where *z* measures the deviation from neutrality. I will demonstrate the effect of this deviation in the results section. Without loss of generality, the extinction rate is set to 1 and only colonization rate and *z* will vary.

### The meta-population model

I then examined the beta diversity-dispersal relationship in the two species meta-population model. The meta-population model assumes that there are infinite number of patches, all patches are equally connected and there is no external immigration so that global extinction is possible. The meta-population model models the occupancy frequency instead of occurrence probability per se, which is also a function of extinction rate *e* and immigration *m*:

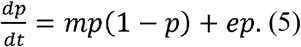

At equilibrium, the occupancy frequency becomes:

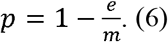

The occurrence probability of a species on any one of the patches equals the occupancy frequency at equilibrium. Assuming equal occurrence probability between two species (*p* = *q*) and independence between islands (e.g. if the two islands are far away from each other), the expected Jaccard dissimilarity is:

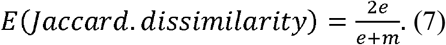

For the non-neutral case, let

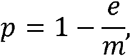

and

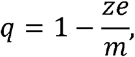

where *z* also measures the deviation from neutrality. The effect of non-neutrality will be shown in the results section. Again, without loss of generality, the extinction rate can be set to 1 and only the migration rate *m* and the deviation from neutrality *z* will be varied.

### The meta-community model

Because the assumptions of independence between species and independence between islands are violated when I take biotic interactions and local dispersal into account, the co-occurrence probability of two species cannot be calculated by the product of marginal occurrence probability of each individual species on each island. Instead, I used a discrete Markov model that built on a previous study (Cazelles *et al.* 2016) to investigate the beta diversity – dispersal relationship. To model co-occurrence patterns, I modeled the state as the whole meta-community occupancy pattern, i.e. a community matrix filled by 0s and 1s with rows representing islands and columns representing species. The transition probability between states are determined by the total number of colonization and extinction events. For example, assuming that at each time step only one event (colonization or no colonization, extinction or no extinction) is allowed for each species on each island, transiting from the 2-species 2-islands metacommunity state of (0,0,1,1) to the state of (1,1,0,0) needs two colonization and two extinction events. A more general formulation is:

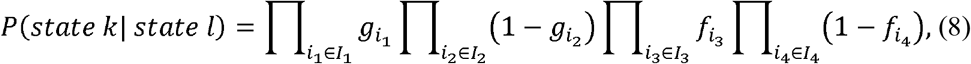

where *g* is the colonization probability, and *f* is the extinction probability. *I*_1_ is the set of colonization events to transit from state *l* to state *k* (transitioning between metacommunity occupancy patterns), *I*_2_ is the set of non-colonization events (an island not colonized by a species at state *l* is still not colonized by the species at state *k*). Similarly, *I*_3_ is the set of extinction events to transit from state *l* to state *k*, *I*_4_ is the set of non-extinction events (a species occupying an island at state *l* does not go extinct at state *k*).

Denote the vector of the probability of initial states (note that each state is an occupancy pattern of the whole meta-community rather than the occurrence probability of each species) as *C*_0_ = [*P*_0_(state 1),…, *P*_0_(state n)]. The stationary distribution of the states can be calculated by iterating the initial state to equilibrium:

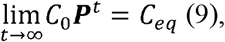

where ****P**** is the transition matrix calculated from equation (8), with element *P_kl_* representing the transition probability from state *l* to state *k*; or by calculating the left eigenvectors of the transition matrix ****P****. The expected Jaccard dissimilarity is then calculated by summing the products of the occurrence probability of every state of the metacommunity and the corresponding Jaccard dissimilarity (Table S1).

The effects of biotic interactions and local dispersals were incorporated by modeling colonization probability *g* as a function of occupancy patterns of the focal species on the neighboring islands, and extinction probability *f* as a function of the occupancy patterns of the interacting species on the same islands. Following Cazelle et al.’s formulation (2016), for species *i* on island *x*, the extinction probability is:

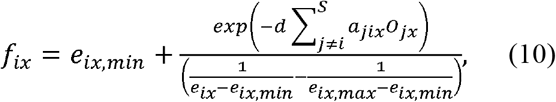

Where *e_ix_* is the baseline extinction probability with no interaction, *e_ix,min_* is the minimum extinction probability (set to 0 throughout this study), *e_ix,max_* is the maximum extinction probability (set to 1 throughout this study), *d* is the total interaction strength, *a_jix_* is the interaction effect of species *j* on species *i*, *O_jx_* is the occurrence of species *j* on island *x* (1 for presence, 0 for absence). *S* is the total number of species in the species pool. The extinction rate *f* is a sigmoid function of the total interaction effect (Appendix, Fig. S1), with positive total interactions lowering the extinction rate of species *i*, negative total interaction increasing the extinction rate of species *i* (Cazelles *et al.* 2016).

To calculate the colonization probability, I combined elements of the island biogeography model and the meta-population model, and I assumed that an empty island *x* can be colonized either by mainland immigrants or immigrants from other occupied islands. I also assumed constant colonization rate from the mainland and that inter-island dispersal is a function of local dispersal rate, island connectivity and island occupancy patterns. Under the assumption that the number of emigrants from an island follows a Poisson distribution and all islands have the same carrying capacity, the probability of having no emigrants from other island is 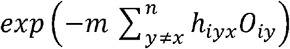, where *m* is the local dispersal(emigration) rate, *h_iyx_* is the connectivity of island *y* to island *x* and *O_iy_* is the occurrence of species *x* on island *y*. The probability of having no immigrants from the mainland is 1 - *c_i_*, where *c_i_*, is the baseline colonization rate (global colonization rate). Then the colonization probability *g_ix_* is the probability of at least one immigrant from the mainland or an immigrant arrives at the focal island:

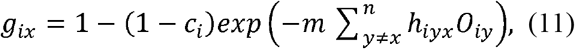

Equation (9) could not be directly applied for large matrices because of large computational costs. Instead of calculating the eigenvector for the transition matrix, I used a Markov chain approach to obtain the stationary distribution of the community matrix (a unique stationary distribution exists for an ergodic Markov chain). The simulation procedure comes naturally from equation (8), where I started with a random community matrix filled with 0s and 1s with rows representing islands and columns representing species (with arbitrary occurrence probability, for example p = 0.2). At each time step, I did the follows:

1. Calculate the extinction probability and colonization probability of each species at each site according to equations (10) and (11).
2. For species *i* at island *x*, if species *i* is present, then it will go extinct with probability *f_ix_* or stay present with probability 1-*f_ix_*; if species *i* is absent, then island *x* will be colonized by species *i* with probability *g_ix_* or stay the same with probability 1-*g_ix_*. The decisions are made by randomly drawing a number from a uniform distribution unif(0,1). If the drawn number is smaller than the given extinction or colonization probability, then species *i* will go extinct or colonize the site *x*; if the drawn number is larger than the given extinction or colonization probability, the occupancy of the site *x* will remain unchanged.
3. Calculate the mean pairwise Jaccard dissimilarity with the new community matrix using the R package ‘betapart’. I also calculated the associated regional richness and mean local richness to aid the visual inspection of equilibrium of metacommunity states.

I iterated the matrix with 1000 steps and take the median mean pairwise Jaccard dissimilarity of the last 100 iterations as the expected mean pairwise Jaccard dissimilarity for the meta-community. Equilibrium for mean pairwise Jaccard dissimilarity, regional richness, mean local richness are generally reached within 1000 iterations (Fig. S2, S3).

#### Parameter settings

##### The two species - two islands competition model

I investigated the beta diversity – local dispersal and beta diversity – global dispersal relationships separately. To highlight the effect of competition, I assumed that the two islands are identical and well connected (*h*_12_ = *h*_21_ = 1, equation 11) and species only differ by baseline extinction rates and interaction effect. To look at the beta diversity – local dispersal relationship, I fixed the global dispersal (baseline dispersal) rate and varied the local dispersal rate. To look at the beta diversity – global dispersal relationship, I fixed the local dispersal rate and varied global dispersal rate. I modeled the effect of competition by setting interactions parameters as *a*_12_ = −5, and *a*_21_ = −3 in equation 10 without loss of generality. The total interaction strength is controlled by *d*, with *d* = 0 indicating no interaction and *d* = 0.1, *d* = 1, *d* = 2 corresponding to different levels of interaction intensity. For the beta diversity – local dispersal relationship, I divided the analysis into four scenarios: 1) low baseline extinction rate and low baseline colonization rate, where *e*_1_ = 0.1, *e*_2_ = 0.2 and *c* =0.1 for both species; 2) low baseline extinction rate and high baseline colonization rate, where *e*_1_ = 0.1, *e*_2_ = 0.2 and *c* = 0.9; 3) high baseline extinction rate and low baseline colonization rate, where *e*_1_ = 0.8, *e*_2_ = 0.9 and *c* =0.1; 4) high baseline extinction rate and high baseline colonization rate, where *e*_1_ = 0.8, *e*_2_ = 0.9 and *c* =0.9. For the beta diversity – global dispersal relationship, I used two levels of global extinction rate (*e*_1_ = 0.1, *e*_2_ = 0.2 for low global extinction rate; *e*_1_ = 0.8, *e*_2_ = 0.9 for high levels of global extinction rate), and three levels of local colonization rate (*m* = 0 for no local dispersal, *m* = 0.1 for low local dispersal and *m* = 10 for high local dispersal). Notice that the baseline colonization rate has to be positive to prevent global extinction in a meta-community simulation with finite number of islands, hence I cannot analyze the equilibrium beta diversity – local dispersal relationship with 0 mainland dispersal rate.

##### The multiple species - multiple islands model

I used 50 species and 9 islands for assessment of the model. Spatial configuration of the 9 islands was laid out as a 3 by 3 grid lattice. The islands are assumed to be identical in their effect on species baseline extinction and baseline colonization (no habitat heterogeneity). The connectivity *h_xy_* among islands in equation (11) is calculated as a negative exponential function of their distances: *h_xy_* = exp[-distance(x,y))] where x and y represents the two-dimensional coordinates of the two islands. The total interaction strength is controlled by *d*, with *d* = 0 indicating no interaction and *d* = 0.01, *d* = 0.1, *d* = 1 corresponding to represent different levels of interaction intensity.

The interaction effect among 50 species were modeled by a random network approach. I first investigated effect of a random network where all interactions are allowed including facilitation, competition and predation. I also explored a competition network in the Appendix (Fig. S6). For the random network model, the interaction coefficients in equation (10) among the 50 species were randomly drawn from a uniform distribution, *a_ij_* ~ unif(−5,5). For beta diversity – local dispersal relationship, I used two levels of global dispersal rate (*c* = 0.001 and *c* = 0.1) and three levels of baseline extinction rates, which is obtained by randomly drawn 50 extinction parameters from a uniform distribution: *e_i_* ~ unif(0.01, 0.02), *e_i_* ~ unif(0.1, 0.2) and *e_i_* ~ unif(0.8, 0.9) respectively. For beta diversity – global dispersal relationship, I used two levels of local dispersal rate (*m* = 0 and *m* =0.1) and the same three levels of baseline extinction rates as described above. For the competition network model, the interaction coefficients in equation (10) among the 50 species were randomly drawn from a uniform distribution, *a_ij_* ~ unif(−5,0). Other parameters are the same with the random network model.

## Results

The relationship between the Jaccard dissimilarity and the occurrence probability of species 1 and 2 (*p* and *q*, respectively) (equation 1) is shown in Figure 1A. If *p* is set to 0 (or close to 0), the Jaccard dissimilarity will decrease monotonically with *q* (Fig. 1A). By contrast, if we assume that the occurrence probability of species *p* is set to 1, then the Jaccard dissimilarity will be a unimodal function of *q* (Fig. 1A). When equally increasing p and q, the Jaccard dissimilarity will decrease monotonically (Fig. 1A). Therefore, the exact shape of the beta diversity – dispersal relationship depends on how dispersal affects the occurrence probabilities of both species.

**Figure 1.**
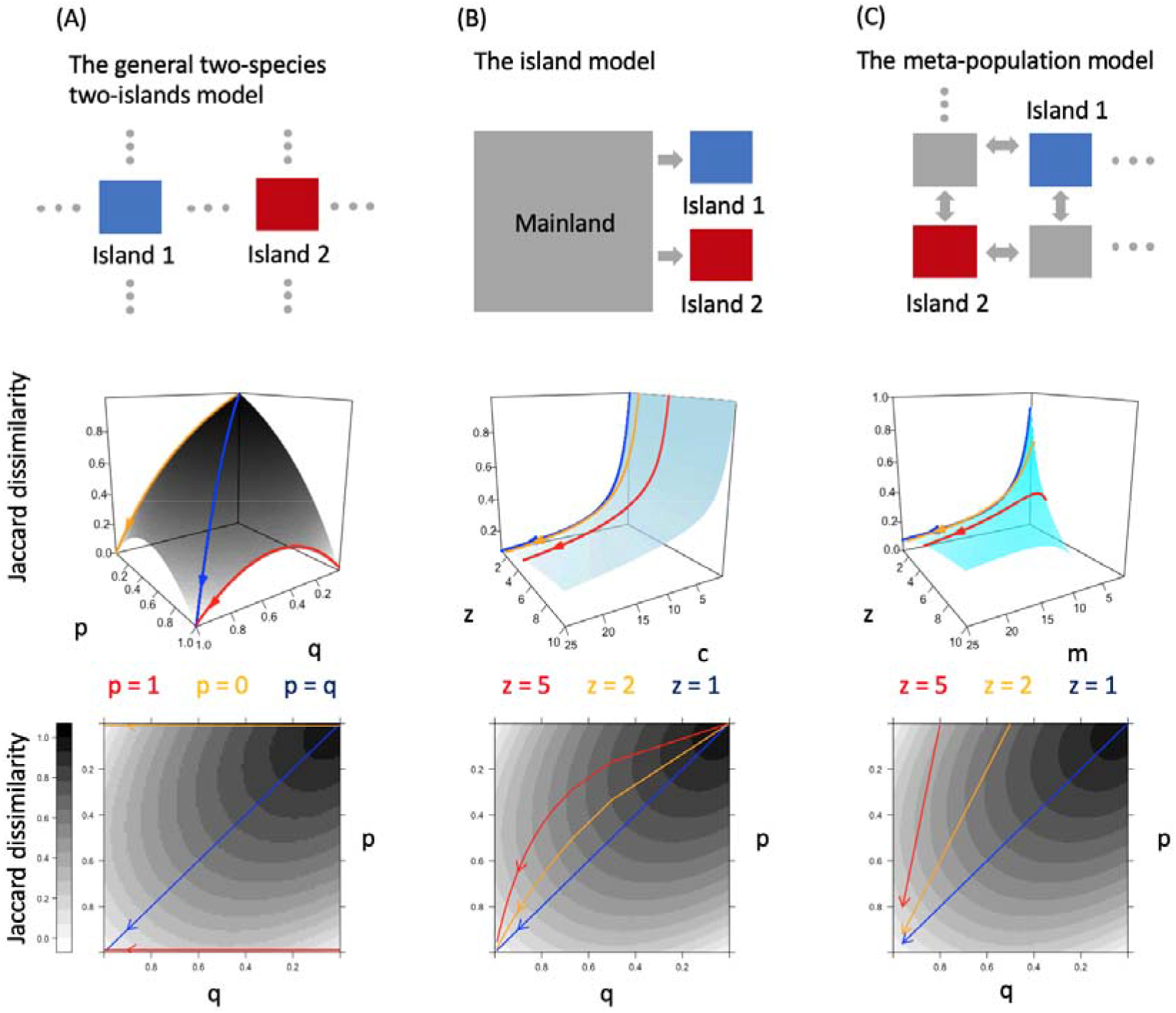
(A) Relationship between Jaccard dissimilarity and the occurrence probability of species 1 (p) and species 2 (q) in the general two-species two-islands model. (B) Relationship between Jaccard dissimilarity and global dispersal rate (c) and the level of non-neutrality (z) in the island model. (C) Relationship between Jaccard dissimilarity and local dispersal rate (m) and the level of non-neutrality (z) in the meta-population model. Arrows in the shaded map indicate the trajectories with increasing dispersal rates for different scenarios on the beta diversity surface of the general two-species two-islands model.

### The island biogeography model

In the neutral two-species island biogeography model (equation 2), the Jaccard dissimilarity is a monotonic decreasing function of the (global) dispersal rate c. When I relaxed the neutral assumption so that the extinction rate of species 2 is *z* times the extinction rate of species 1, the Jaccard dissimilarity remains a monotonic decreasing function of (global) dispersal rate *c* (Fig. 1B).

### The meta-population model

In the neutral two-species meta-population model (equation 3), Jaccard dissimilarity is also a monotonic decreasing function of (local) dispersal rate *m*. But when I increased the level of non-neutrality, characterized by the parameter *z*, the Jaccard dissimilarity becomes a unimodal function of local dispersal rate (Fig. 1C).

### The meta-community model

In the low baseline extinction rate and low baseline (global) colonization rate scenario, Jaccard dissimilarity decreases with local dispersal rate when there is no competition. When the competition is weak, Jaccard dissimilarity is higher than the case of no interaction, but the beta diversity – dispersal relationship is still a monotonic decreasing function. When total competition is strong, Jaccard dissimilarity first decreases and then increases with the local dispersal rate (Fig. 2A). The equilibrium occurrence probabilities of the two species increase with local dispersal when total interaction strength is high, but occurrence probability of species 1 (the superior competitor) increases faster with higher dispersal than species 2. In the low baseline extinction rate, high baseline (global) colonization rate scenario, increasing total competition also changes the beta diversity – local dispersal relationship from a monotonic decreasing function to an increasing function. The equilibrium occurrence probabilities of the two species also increase with local dispersal rate. In the high baseline extinction, low baseline colonization scenario, the Jaccard dissimilarity first decreases then increases with local dispersal rate with or without competition (Fig. 2C). In the high baseline extinction, high baseline colonization scenario, the Jaccard dissimilarity increases with local dispersal rate in all levels of competition strength. But stronger competition decreases the overall beta diversity (Fig. 2D).

**Figure 2.**
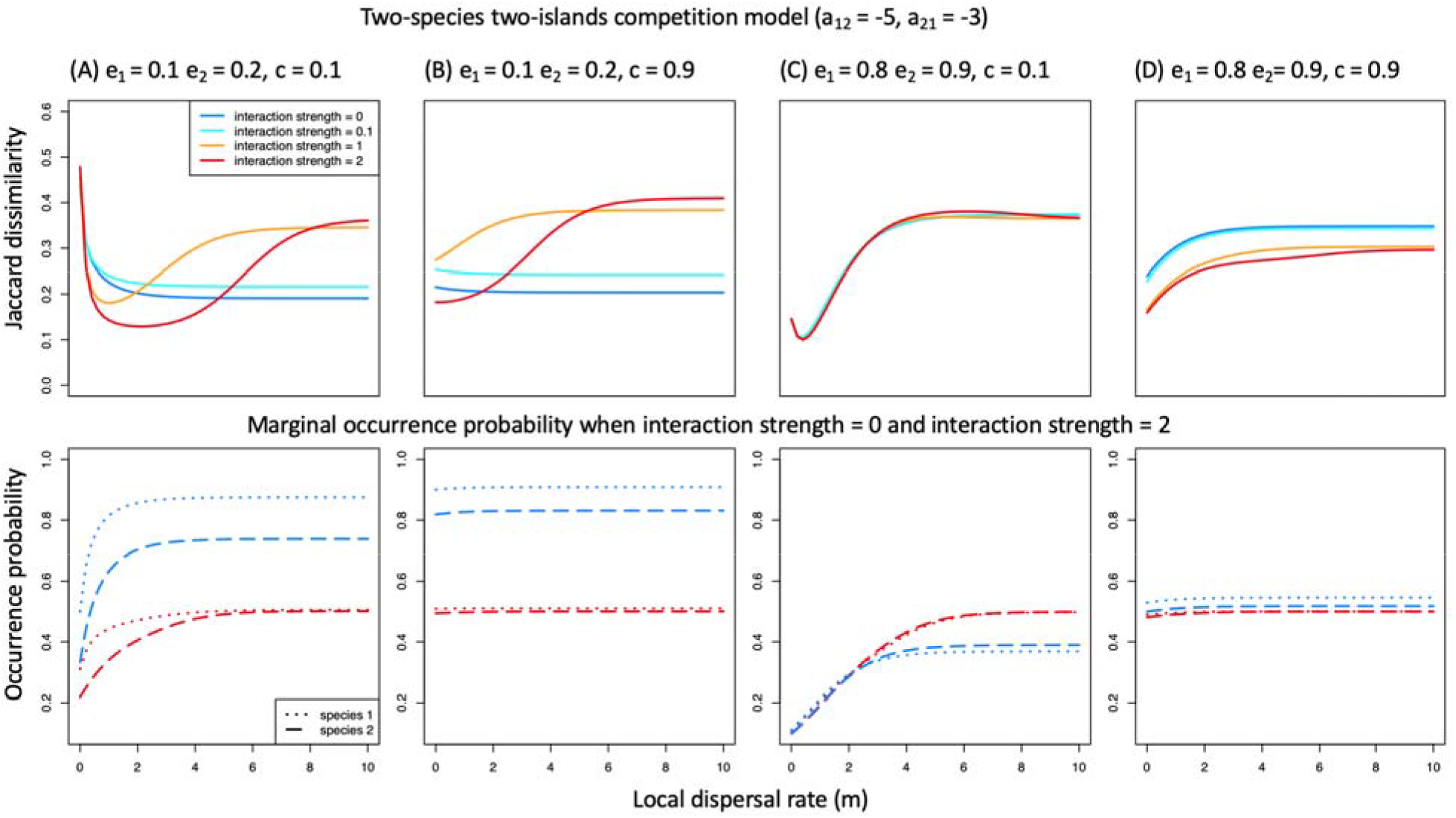
Relationship between Jaccard dissimilarity and local dispersal rate in the two-species two-islands metacommunity model when global colonization rate is fixed in (A) the low baseline extinction, low global colonization scenario; (B) the low baseline extinction, high global colonization scenario; (C) the high baseline extinction, low global colonization scenario; (D) the high baseline extinction, high global colonization scenario. The red lines in the bottom panels correspond to the equilibrium occurrence probabilities of the two species when interaction strength = 2.

Then I fixed the local dispersal rate and looked at the beta diversity – global dispersal relationship. When there is no local dispersal and the baseline extinction rates are low and competition is weak, the Jaccard dissimilarity decreases monotonically with global dispersal (Fig 3A). By contrast, when competition is strong, the Jaccard dissimilarity exhibits a non-monotonic relationship with global dispersal, increasing at first and then decreasing with it (Fig. 3A). When there is no local dispersal and the baseline extinction rates are high, the Jaccard dissimilarity decreases with global dispersal for all levels of competition strength (Fig. 3B). With the presence of local dispersal, the Jaccard dissimilarity becomes a unimodal function of global dispersal for all levels of baseline extinction rates, local dispersal rate and competition strength (Fig. 3C-F). Higher local dispersal rate leads to higher maximum Jaccard dissimilarity when baseline extinction rate is high (Fig. 3F).

**Figure 3.**
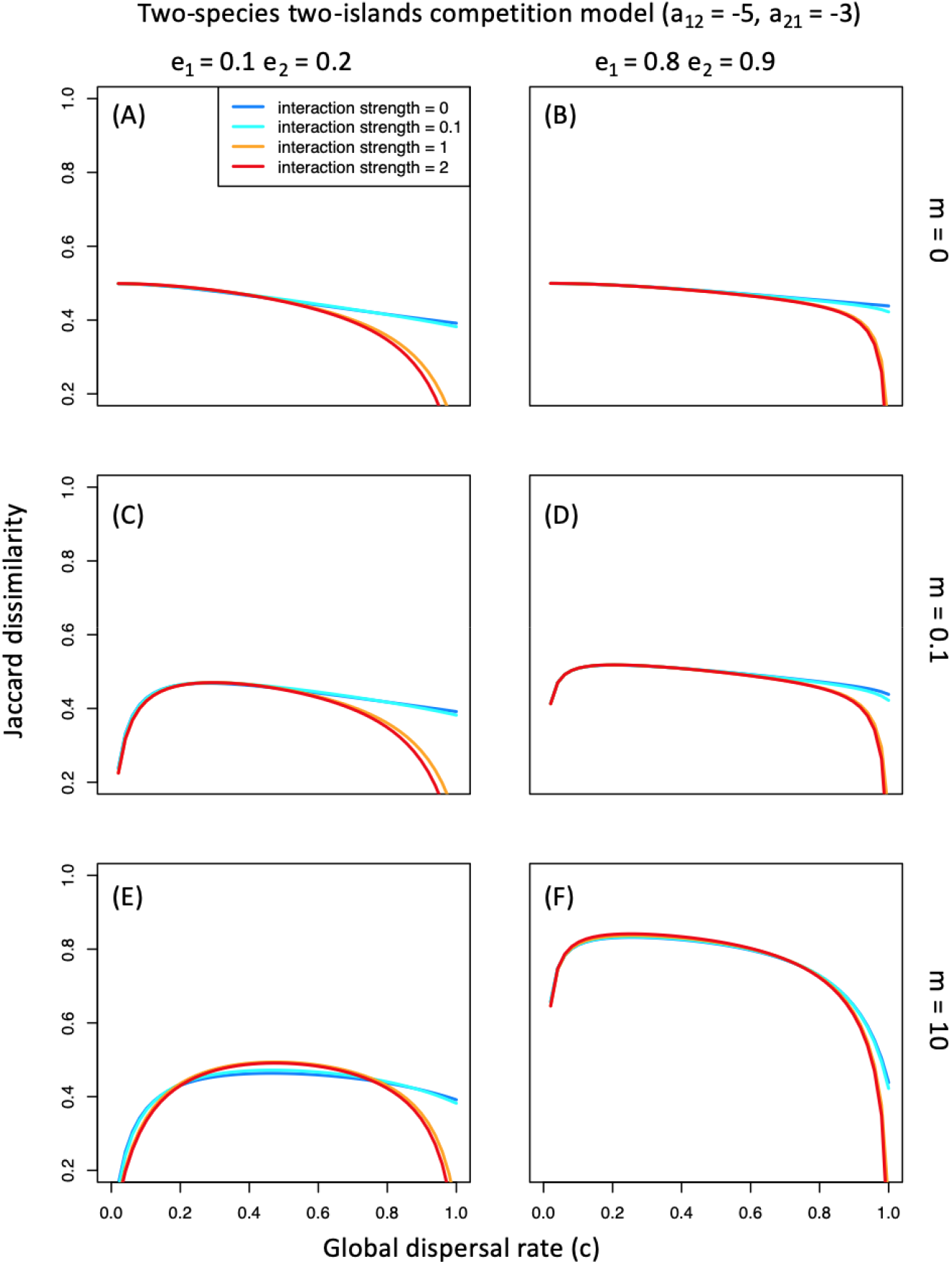
Relationship between Jaccard dissimilarity and global dispersal rate in the two-species two-islands metacommunity model when local dispersal rate is fixed. (A) low baseline extinction, no local dispersal; (B) high baseline extinction, no local dispersal; (C) low baseline extinction, low local dispersal; (D) high baseline extinction, low local dispersal; (E) low baseline extinction, high local dispersal; (F) high baseline extinction, high local dispersal.

### Multiple species – multiple islands model: random network

For the 50 species - 9 islands random network model, when baseline extinction rates and baseline colonization rates are low, increasing interaction strength changes the beta diversity – local dispersal relationship from a decreasing function to an increasing function of local dispersal rate (Fig. 4A, B). When baseline extinction rates are high and baseline colonization rates are low, the Jaccard dissimilarity is a unimodal function of local dispersal at all levels of biotic interactions (Fig. 5C). When baseline colonization rates are high, the Jaccard dissimilarity only decreases monotonically with local dispersal rate (Fig. 4D-F).

**Figure 4.**
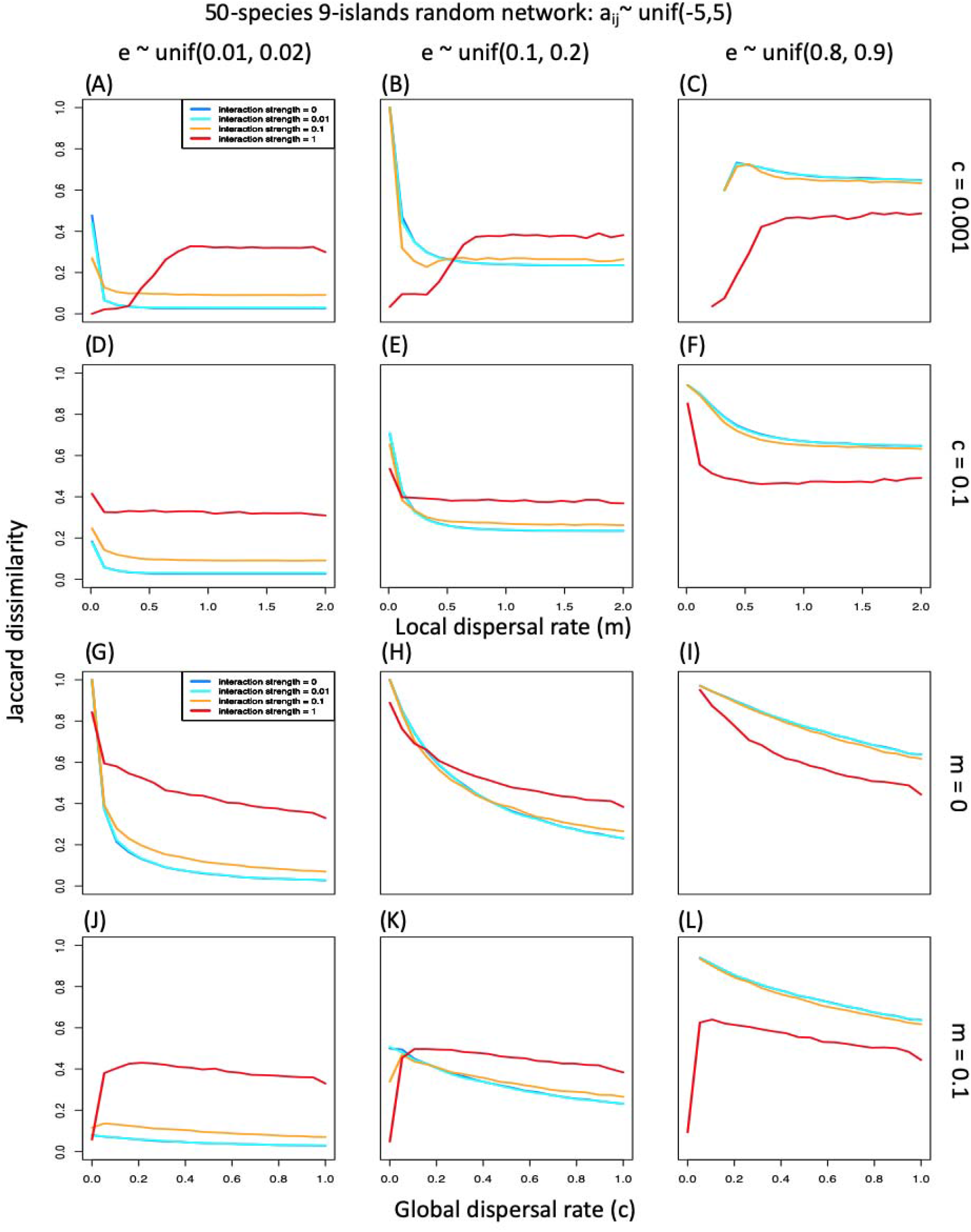
(A) – (F) Relationship between Jaccard dissimilarity and local dispersal rate in the 50-species 9-islands metacommunity model with random network a_ij_ ~unif(−5,5). (G) – (L) Relationship between Jaccard dissimilarity and global dispersal rate in the 50-species 9-islands metacommunity model with random network.

When there is no local dispersal, the Jaccard dissimilarity decreases with global dispersal for all levels of the baseline extinction rate with all levels of biotic interactions (Fig. 4G-I). When local dispersal is present, increasing interaction strength changes the beta diversity – global dispersal relationship from a decreasing function to an increasing function of the global dispersal rate (Fig. 4J-L).

## Discussion

### Theoretical results

Using a simple meta-community model built on the classic island biogeography model and meta-population model, I show that beta diversity exhibits a rich range of responses to the dispersal rate, ranging from a monotonic decrease to a ‘U’ shaped curve or a hump shaped curve. The shape of the beta diversity – dispersal relationship depends on the type of dispersal (global dispersal or local dispersal), the strength of biotic interactions, and the rates of baseline extinction and colonization. Here, I will discuss the theoretical results and how they could guide us in further empirical studies.

In essence, the increasing beta diversity – dispersal relationship reflects variations in species-level occurrence probability – dispersal relationship. To understand how this could happen, let’s consider two hypothetical scenarios where the occurrence probability of species 1 is fixed near 0 or 1 (Fig. 1A). In the former case, beta diversity decreases monotonically with dispersal rate because increasing occurrence probability of species 2 will only have a homogenizing effect on the two-islands meta-community as more islands gets occupied by a single species 2; in the latter case, beta diversity will first increase with dispersal rate because increasing occurrence probability of species 2 breaks the dominance of species 1 across islands (imagine we are randomly drawing two islands from a bunch of islands for comparison) at the intermediate dispersal rate, but will eventually decrease because more islands will be saturated with both species. To explain this in relation to gamma and alpha diversity: whenever gamma diversity (of any two island pairs) increases faster than local richness with higher dispersal rate, we observe an increasing beta diversity – dispersal relationship.

For both the island biogeography model and the meta-population model, assuming neutrality produces a monotonicly decreasing relationship between beta diversity and dispersal. When they deviate from the neutral scenario, the beta diversity – dispersal relationship in the island biogeography model remains a monotonicly decreasing function, but in the meta-population model shifts to hump shaped curves with higher level of non-neutrality. The difference between the models arises from the fact that dispersal rate of the meta-population model depends on occupancy frequency which creates more variation in species-level occurrence probability-dispersal relationship and makes it more likely to traverse the unimodal surface of beta diversity in the general two-species two-islands model when I add variations in parameters (Fig. 1B, C). This might explain why the hump shaped beta diversity – dispersal relationship is not detected in previous theoretical work, which had focused on the neutral scenario (Hubbell 2001; Mouquet & Loreau 2003; Lu *et al.* 2019).

In a two-species two-islands meta-community model with species competition. At low baseline extinction and low baseline colonization (global dispersal) rates, increasing competition changes the beta diversity-dispersal relationship from monotonicly decreasing to ‘U’ shaped. The initial decrease of Jaccard dissimilarity with dispersal rate results from the fact that when local dispersal rate is low, the model behaves like an island biogeography model where the islands are relatively independent. The later increase of Jaccard dissimilarity with dispersal is the signal of enhanced species competition when islands become more connected and local co-occurrence becomes less likely. In the low baseline extinction and high baseline colonization (global dispersal) rates scenario, the early increase of local dispersal does not lead to initial decrease of Jaccard dissimilarity because the baseline occurrence probabilities of both species are high (Fig. 2B). In the two high baseline extinction rate scenarios, competition does not significantly change the shape of the beta diversity – dispersal relationship (Fig. 3D, F) because competition only changes the extinction rates by a small amount (notice that the maximum extinction probability is bounded by 1).

Fixing the local dispersal rate and investigating the beta diversity – global dispersal relationship gives us a better appreciation of the difference between these two types of dispersal: in the absence of local dispersal and biotic interactions, global dispersal has a consistent homogenizing effect on local communities (Fig. 3A, B). A hump-shaped curve only emerges when local dispersal is non-vanishing (Fig. 3C-F). In summary, local dispersal plays a dominant role in giving rise to the hump shaped beta diversity – dispersal relationship in competition models. This message is demonstrated clearer by the 50 species random model (Fig. 4G-I) as well as the competition model (Fig. S6G-I) where global dispersal only leads to homogenization when there is no local dispersal, and that the hump shaped beta diversity – local dispersal relationship could be masked by high global dispersal (Fig. 4D-F). Moreover, biotic interaction creates more variation in species-level occurrence probability – dispersal relationships which contributes to the increase of beta diversity (Fig. S4, S5). The results of the 50 species competition network model are qualitatively consistent with the random network model (Fig. S6).

### Implications for empirical studies

There are a few reasons why the phenomenon that dispersal enhances beta diversity is overlooked before. For example, as suggested by Vannette and Fukami (2017), the neglection of biotic interactions and the unrealistic manipulation of dispersal by bulk propagule addition etc. I offer a few considerations here. First, the level of stochasticity in dispersal (as well as extinction) differs between studies. Because the results presented in this paper are derived from a stochastic occupancy model, an experiment that conducted on grassland and manipulated the amounts of seed dispersal might not detect what I found here because the processes operate mostly on abundance variation (Ron *et al.* 2018). Many microcosm experiments also did not implement the dispersal process in a stochastic manner, which might prevent the observation of hump shaped or positive beta diversity – dispersal relationship (Grainger & Gilbert 2016). Second, it has rarely been emphasized that global dispersal and local dispersal might have different effects on beta community patterns. The distinction between global dispersal and local dispersal dictates the method to manipulate dispersal. For example, the commonly used ‘merge and redistribute’ technique mimics global dispersal by forcing all patches to share a well-mixed and relatively unchanged species pool (Grainger & Gilbert 2016); on the contrary, Vannette and Fukami’s bagged procedure manipulated local dispersal by limiting the access of pollinators (2017). While the global dispersal experiments are abundant, the local dispersal experiments are scarce (Grainger & Gilbert 2016). Third, the relative amount of global and local dispersal also differs between studied systems. In the nectar microbe experiment, the system might be similar to a meta-community model with limited external dispersal given the large number of patches (flowers in Vannette and Fukami’s case). But in a 25-hectare forest plot, external dispersal might not be negligible especially when the dispersal kernel for tree species has a heavy tail (Clark 1998). The microcosms have the advantage of having a closed system but, as mentioned above, the distinction between local dispersal and global dispersal has seldom been emphasized. Finally, the details of biotic interactions should be fully considered. A tropical forest plot might be dominated by diffuse competition among tree species while zooplankton communities could include multiple trophic levels.

## Conclusions

Using a metacommunity model, I shed light on the many possible relationships between beta-diversity and dispersal, and their mechanistic underpinnings. Stochasticity, non-neutrality, dispersal types, and biotic interactions all play prominent roles in determining the specific relationships observed. My results motivate new experiments where the dispersal type, and the closeness of the studied system can be manipulated to provide richer insights on meta-community dynamics.

## Acknowledgements

Biggest thank to Alvaro Sanchez who offered tremendous help and feedback throughout the development of this project and the writing. Thank David Vasseur for providing feedbacks in the early development of the work. Thank Jussi Mäkinen and Charlie Marsh for comments on the first draft.

