## Supplementary material for "Letters: Complex response of beta diversity to dispersal in meta-community models": Table S1

Supplements

Beta diversity – dispersal relationship

Muyang Lu

**Derivation of the two-species two-islands model:**

Table S1. All possible states of a two-species two-islands metacommunity and the associated probabilities and beta diversity values by assuming independence among islands and species.

| state | species1, island1 | species2, island1 | species1, island2 | species2, island2 | Probability | Jaccard dissimilarity |
| --- | --- | --- | --- | --- | --- | --- |
| 1 | 0 | 0 | 0 | 0 | (1-p)^2^(1-q)^2^ | NAN |
| 2 | 1 | 0 | 0 | 0 | p(1-p)(1-q)^2^ | 1 |
| 3 | 0 | 1 | 0 | 0 | p(1-p)(1-q)^2^ | 1 |
| 4 | 1 | 1 | 0 | 0 | p^2^(1-q)^2^ | 1 |
| 5 | 0 | 0 | 1 | 0 | (1-p)^2^(1-q)q | 1 |
| 6 | 1 | 0 | 1 | 0 | p(1-p)q(1-q) | 0 |
| 7 | 0 | 1 | 1 | 0 | p(1-p)q(1-q) | 1 |
| 8 | 1 | 1 | 1 | 0 | p^2^(1-q)q | 0.5 |
| 9 | 0 | 0 | 0 | 1 | (1-p)^2^(1-q)q | 1 |
| 10 | 1 | 0 | 0 | 1 | p(1-p)(1-q)^2^ | 1 |
| 11 | 0 | 1 | 0 | 1 | p(1-p)q(1-q) | 0 |
| 12 | 1 | 1 | 0 | 1 | p^2^(1-q)q | 0.5 |
| 13 | 0 | 0 | 1 | 1 | (1-p)^2^q^2^ | 1 |
| 14 | 1 | 0 | 1 | 1 | p(1-p)q^2^ | 0.5 |
| 15 | 0 | 1 | 1 | 1 | p(1-p)q^2^ | 0.5 |
| 16 | 1 | 1 | 1 | 1 | p^2^q2 | 0 |

The expected Jaccard dissimilarity is calculated by summing the product of probability and the Jaccard dissimilarity of each state and renormalized by 1-(1-p)^2^(1-q)^2^ because double-absence is excluded in calculating beta diversity.

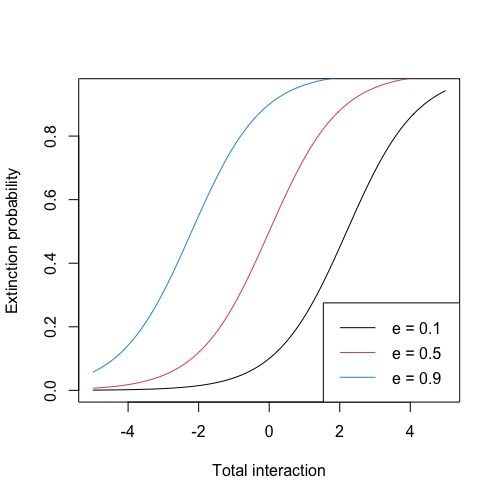

Figure S1. Extinction probability as function of the baseline extinction probability e.

**Examples of convergence plots when interaction strength = 1 for the 50 species – 9 islands competition model, c = 0.0001, e ~ unif(0.01,0.02), P0 = 0.2:**

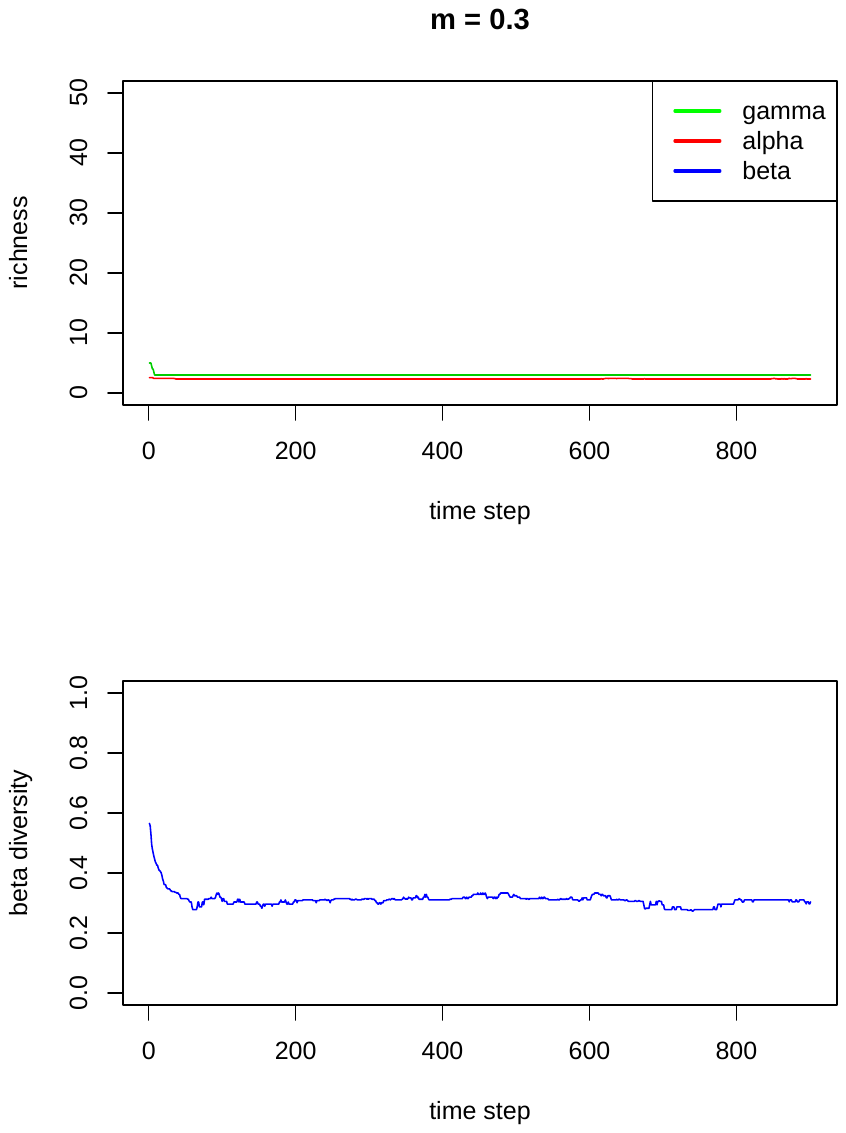

Figure S2. Gamma, alpha and beta diversity plotted against number of iterations when m = 0.3.

Table S2. Occurrence probability estimated as the mean occurrence of the last 100 iterations when m = 0.3. Species not shown have 0 occurrence.

| species | island.1 | island.2 | island.3 | island.4 | island.5 | island.6 | island.7 | island.8 | island.9 |
| --- | --- | --- | --- | --- | --- | --- | --- | --- | --- |
| 22 | 0.97 | 0.98 | 0.95 | 0.94 | 0.96 | 0.94 | 0.96 | 0.89 | 0.96 |
| 28 | 0.77 | 0.84 | 0.84 | 0.87 | 0.87 | 0.8 | 0.84 | 0.84 | 0.67 |
| 7 | 0.6 | 0.59 | 0.65 | 0.51 | 0.65 | 0.72 | 0.45 | 0.62 | 0.6 |

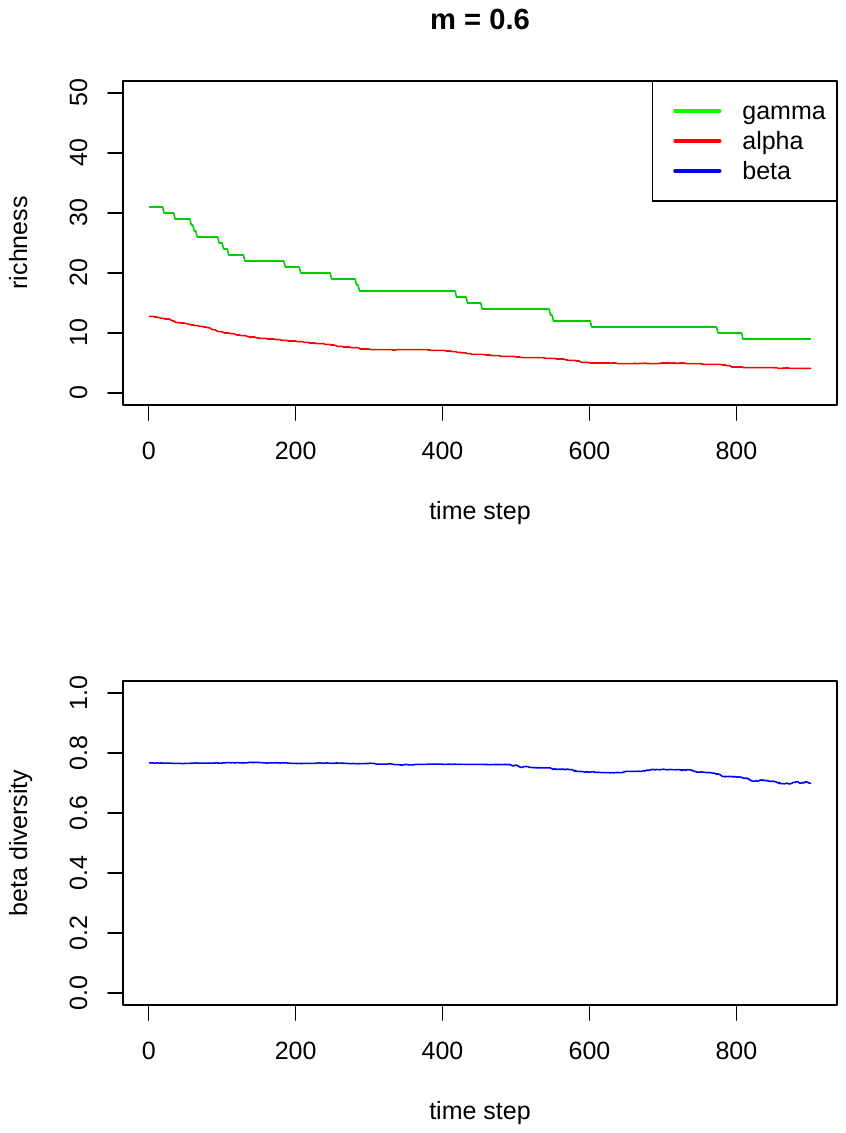

Figure S3. Gamma, alpha and beta diversity plotted against number of iterations when m = 0.6.

Table S3. Occurrence probability estimated as the mean occurrence of the last 100 iterations when m = 0.6. Species not shown have 0 occurrence.

| species | island.1 | island.2 | island.3 | island.4 | island.5 | island.6 | island.7 | island.8 | island.9 |
| --- | --- | --- | --- | --- | --- | --- | --- | --- | --- |
| 23 | 0.49 | 0.45 | 0.44 | 0.47 | 0.48 | 0.46 | 0.48 | 0.46 | 0.44 |
| 9 | 0.48 | 0.49 | 0.42 | 0.49 | 0.48 | 0.45 | 0.45 | 0.45 | 0.48 |
| 37 | 0.47 | 0.44 | 0.49 | 0.45 | 0.5 | 0.48 | 0.48 | 0.49 | 0.47 |
| 19 | 0.46 | 0.47 | 0.43 | 0.45 | 0.48 | 0.46 | 0.38 | 0.44 | 0.41 |
| 33 | 0.46 | 0.48 | 0.47 | 0.47 | 0.45 | 0.45 | 0.4 | 0.43 | 0.42 |
| 44 | 0.43 | 0.44 | 0.45 | 0.46 | 0.5 | 0.48 | 0.5 | 0.46 | 0.46 |
| 11 | 0.42 | 0.46 | 0.41 | 0.48 | 0.49 | 0.46 | 0.42 | 0.45 | 0.47 |
| 14 | 0.41 | 0.44 | 0.48 | 0.44 | 0.48 | 0.49 | 0.49 | 0.47 | 0.46 |
| 17 | 0.41 | 0.45 | 0.45 | 0.45 | 0.5 | 0.48 | 0.4 | 0.47 | 0.41 |

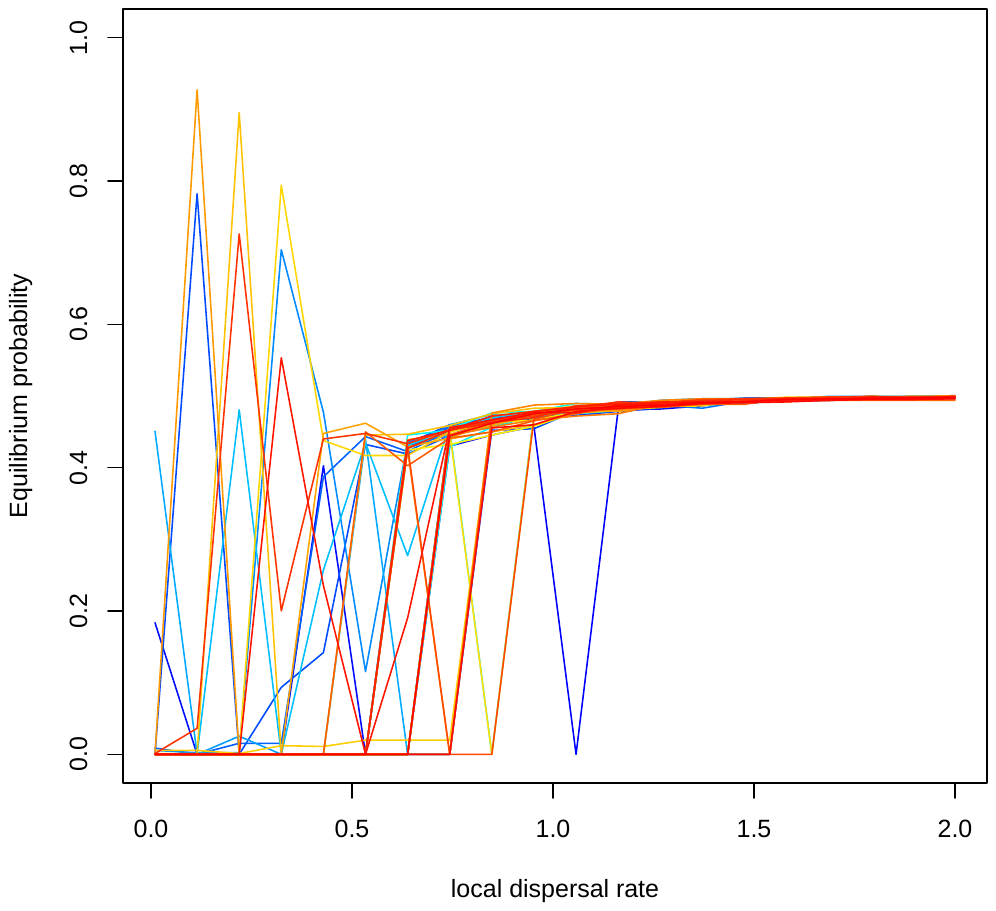

Figure S4. Mean equilibrium occurrence probability for 50 species plotted against local dispersal rate.

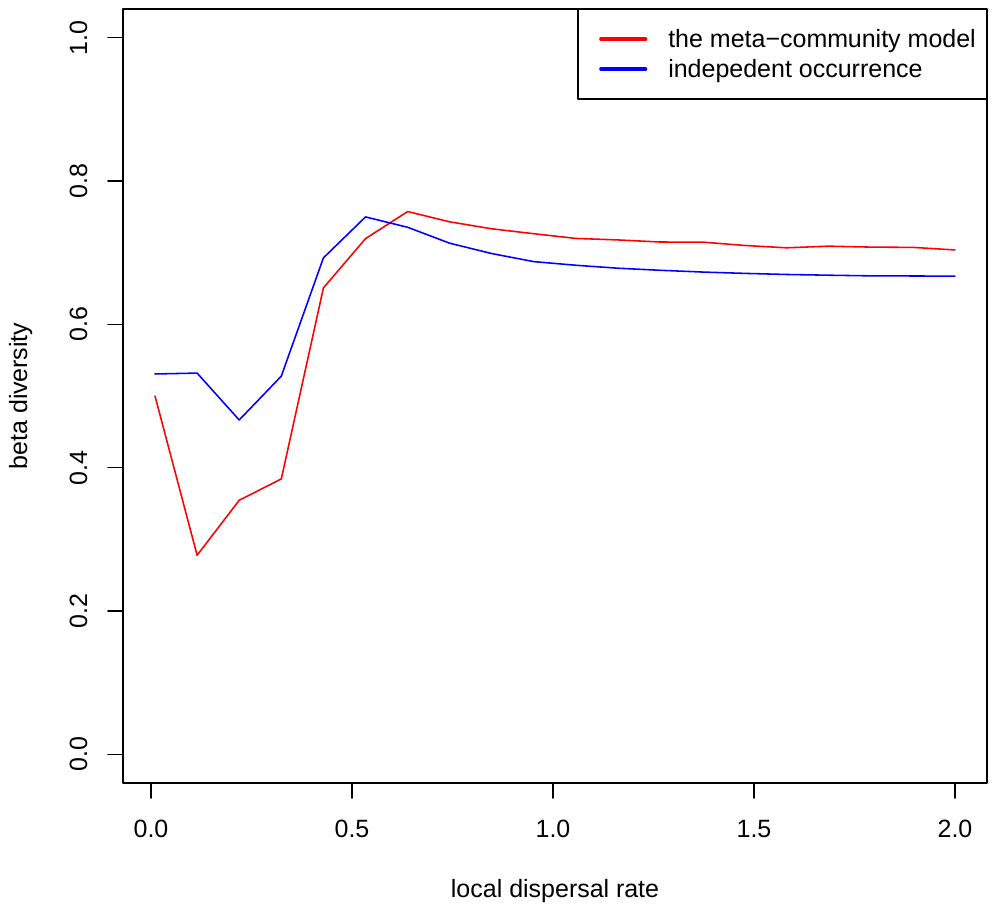

Figure S5. Beta diversity calculated from the simulated meta-community model compared to beta diversity estimated by assuming independent occurrence probability among species and islands, suggesting the humped-shape relationship solely arise from species-level variation in occurrence probability-dispersal relationship.

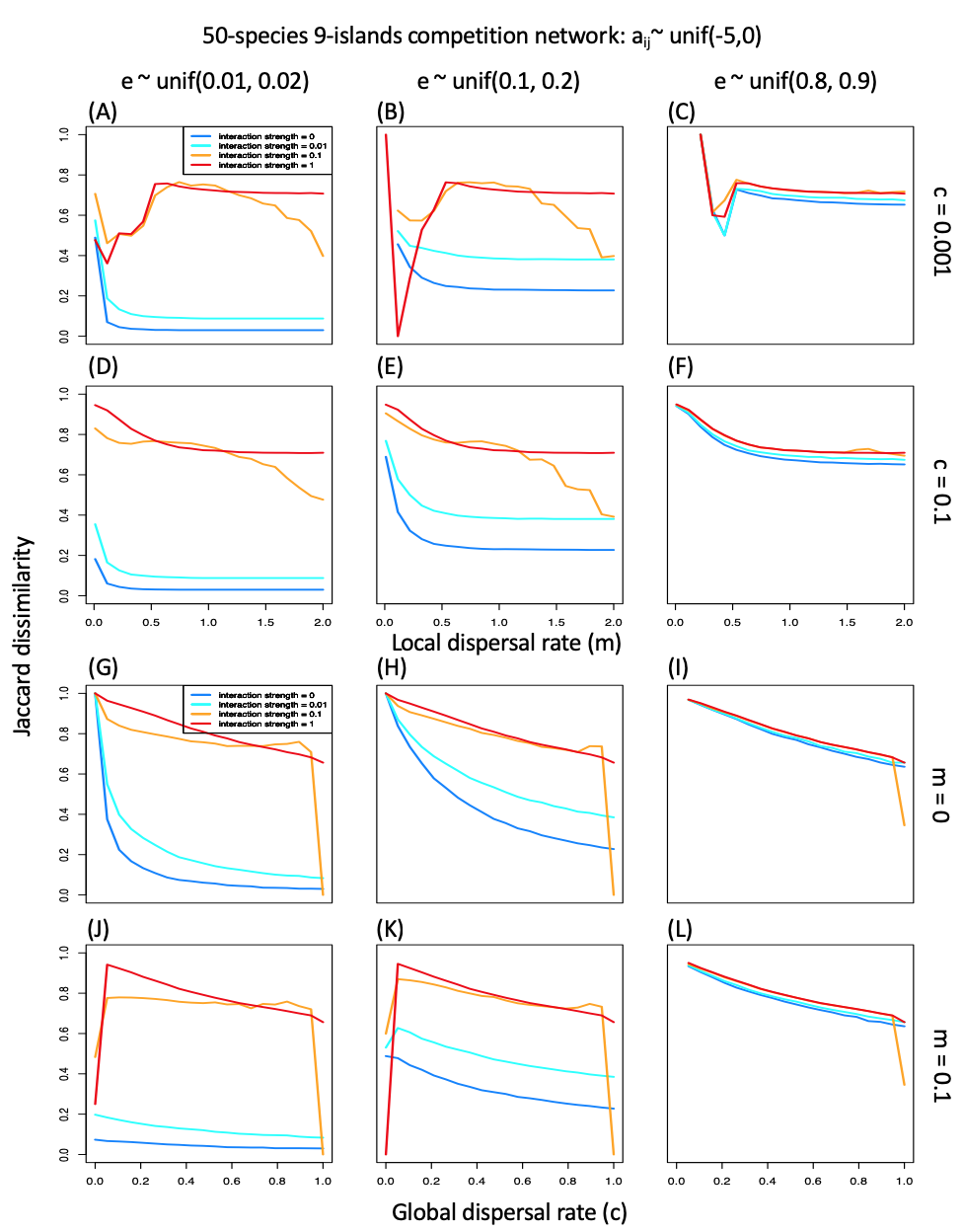

Fig. S6 (A) – (F) Relationship between Jaccard dissimilarity and local dispersal rate in the 50-species 9-islands metacommunity model with competition network a_ij_ ~unif(-5,0). (G) – (L) Relationship between Jaccard dissimilarity and global dispersal rate in the 50-species 9-islands metacommunity model with competition network.
